# Genetic and phenotypic analysis of the virulence plasmid of a non-Shigatoxigenic enteroaggregative *Escherichia coli* O104:H4 outbreak strain

**DOI:** 10.1101/2024.10.31.621240

**Authors:** Rachel Whelan, Martyna Cyganek, Benjamin Dickins, Jonathan C Thomas, Gareth McVicker

## Abstract

Enteroaggregative *Escherichia coli* O104:H4 is best known for causing a worldwide outbreak in 2011 due to acquisition of a Shiga-like toxin alongside traditional enteroaggregative virulence traits; however, whilst the 2011 outbreak strain has been well-studied, the virulence plasmid of O104:H4 has been subjected to far less experimental analysis. In this paper, we analyse the genetic and phenotypic contribution of the pAA virulence plasmid to a non-Shigatoxigenic O104:H4 strain (1070/13) that was nonetheless implicated in a substantial UK outbreak in 2013. We find that pAA_1070_ is 99.95% identical across 88% of the plasmid sequence to pTY2 from the 2011 outbreak strain and has a copy number of approximately 2-3 plasmid molecules per chromosome. We demonstrate that pAA_1070_ carries a functional CcdAB plasmid addiction system that only marginally impacts its stability under the conditions tested and that none of the other toxin-antitoxin systems encoded by the plasmid appear to be functional, though we note a surprisingly high stability of the plasmid *in vitro* regardless. We demonstrate the expected contribution of pAA_1070_ to intestinal cell adhesion but find that it does not contribute to biofilm formation. When assessing the impact of pAA_1070_ on motility, we discovered a region of the O104:H4 chromosome that can be excised, abolishing motility via truncation of the *fliR* gene. Ultimately this work contributes to our knowledge of enteroaggregative *E. coli* as an important pathovar in its own right and demonstrates the complexity but necessity of experimentally characterising genuine outbreak strains rather than domesticated laboratory strains in order to understand virulence phenotypes.

**IMPACT STATEMENT:** Enteroaggregative *Escherichia coli* causes severe food-borne illness in humans. One serotype, O104:H4, caused a worldwide outbreak in 2011 that infected almost 4,000 people and killed dozens. The 2011 strain contained two key virulence factors: the Shiga toxin encoded by a bacteriophage and a plasmid that encodes aggregative adherence fimbriae for adhesion to host cell surfaces. Whilst the Shiga toxin has been studied extensively, the virulence plasmid has been far less well-characterised. In this study, we characterise the plasmid of a non-Shigatoxigenic O104:H4 outbreak strain and find that it contributes to host cell adherence, as expected, and that it encodes a functional CcdAB toxin-antitoxin stability system. We also demonstrate that part of the strain’s chromosome can be spontaneously lost during growth in the laboratory, converting the organism from highly-motile to non-motile, and suggesting a future evolutionary route for the bacterium, as has occurred in similar enteric pathogens such as *Shigella*. In summary, this work contributes to our knowledge of often-neglected enteroaggregative *E. coli* as an important pathovar and highlights the necessity of experimentally characterising genuine outbreak strains rather than “domesticated” laboratory strains.

**Repositories/Data Summary:** The authors confirm all supporting data, code and protocols have been provided within the article or through supplementary data files. Hybrid whole-genome sequence data is available for strains 1070/13 (accession numbers CP171474, CP171475 and CP171476) and 1070/13 pAA^−^ (accession numbers CP171472 and CP171473). Sequencing data for 1070/13 produced by Dallman *et al.* (2014) is available from the European Nucleotide Archive: sample accession SAMN02730196.

## INTRODUCTION

*Escherichia coli* is a common enteric bacterium in mammals. Due to its propensity for mobile genetic element (MGE) acquisition, commensal or otherwise avirulent *E. coli* can be converted into pathogenic forms known as pathovars, defined by their mode of virulence in human disease [1]. One such diarrheagenic pathovar, enteroaggregative *E. coli* (EAEC), is phenotypically characterised by a “stacked-brick” adherence pattern on intestinal epithelial cells [2]. This phenotype is thought to result from the production of aggregative adherence fimbriae (AAF) typically encoded on a large (70-130 kb) aggregative adherence plasmid, pAA.

In mid-2011, EAEC serotype O104:H4 made headlines after infecting almost 4,000 people and killing over 50 otherwise-healthy adults [3]. The atypical outbreak variant was found to be a hybrid pathovar, carrying both the expected EAEC virulence genes on its pAA variant, named pTY2 in sequenced strain TY2482 [4,5], and a gene encoding Shiga-like toxin variant Stx2a on a prophage [6,3]. The carriage of both traits likely dramatically enhanced the strain’s virulence relative to both non-enteroaggregative enterohaemorrhagic *E. coli* (EHEC) and non-enterohaemorrhagic EAEC. Whilst Shiga-like toxins are highly-studied virulence factors of EHEC, the virulence plasmid of EAEC is far less well-characterised. AAF are found in five different forms [7–11] and the pAA plasmids that encode them are similarly varied. After a thorough recent analysis, Boisen *et al.* [12] proposed that EAEC be characterised genetically by the presence of both AAF and the plasmid-encoded master regulator, AggR. Other genes such as those encoding the anti-aggregation protein dispersin and its related transporters are also highly-represented amongst EAEC strains [13–16].

Given their size, low copy number and propensity to encode complex protein-based macromolecules (e.g. specialised fimbriae or secretion systems), enteric virulence plasmids are often associated with a growth burden and can be lost or degraded when their specific function is not required, such as during growth outside the pathogen’s preferred niche or in laboratory culture. To counter this, many such plasmids encode maintenance systems or induce compensatory mutations in the bacterial host [17,18]. Toxin-antitoxin (TA) loci are plasmid maintenance systems comprised of genes encoding a stable toxin protein and unstable antitoxin, which can itself be either a protein or RNA molecule. In type II TA systems, both toxin and antitoxin are proteins, and the antitoxin functions by directly binding to and sequestering the toxin [19]. Type II antitoxins are rapidly degraded by cellular proteases such as Lon [20,21]. After bacterial cell division, loss of a plasmid carrying TA genes results in toxin activation and bacterial growth arrest or death, as the antitoxin is degraded and not replaced; hence, a growing population must retain the plasmid to avoid this post-segregational killing (PSK). Two well-characterised type II TA systems are *ccdAB*, encoding DNA gyrase poison CcdB and its antidote CcdA [22], and *vapBC*, encoding tRNA-cleaving toxin VapC and its antidote VapB [23,21]. Both *ccdAB* and *vapBC* are common in *E. coli* and *Shigella* virulence plasmids, though their stability functions vary [24–27].

In 2013, a multi-pathogen foodborne disease outbreak in Newcastle, United Kingdom [28] was found to be predominantly caused by EAEC strains carrying a wide range of pAA plasmids. Whilst these isolates were all Shiga toxin-negative, amongst the strains were O104:H4 isolates carrying a form of pAA almost identical to pTY2 from the 2011 outbreak [28]. The 2013 Newcastle strains can therefore be used as convenient model organisms to study the virulence and maintenance functions of this particular pAA variant.

Given their role in pathoadaption, knowledge of the impact and stability of *E. coli* MGEs is essential to understand how the organism causes disease and how it is able to amass multiple MGEs in order to form hybrid pathovars. In this study, we characterise various virulence functions of pAA in Newcastle outbreak strain 1070/13 and analyse the role of CcdAB encoded on this plasmid. We also identify a region within the chromosome of the organism that is able to spontaneously excise and disable motility in the strain, perhaps suggesting a future evolutionary step for the organism.

## METHODS

### Bacterial strains and growth conditions

Bacterial strains (**Supplementary Table 1**) were grown in lysogeny broth (LB; Sigma-Aldrich, UK) or on solid L-agar (LB + 1.5 % (w/v) agar (Sigma-Aldrich, UK)) supplemented with relevant antibiotics unless otherwise stated. Bacteria were incubated at 37 °C (with aeration for liquid cultures) unless otherwise stated. Antibiotics (Sigma-Aldrich, UK) were used at the following final concentrations for selection and plasmid maintenance: neomycin 50 µg/ml; chloramphenicol 30 µg/ml; ampicillin 100 µg/ml. Sucrose agar for quantification of pSTAB plasmid loss was made according to the following recipe: 10 g/L tryptone (Sigma-Aldrich, UK), 5 g/L yeast extract (Fisher Scientific, UK), 10 % (w/v) sucrose (Sigma-Aldrich, UK) and 1.5 % (w/v) agar. Where required for growth of auxotrophs, diaminopimelic acid (DAP; Sigma-Aldrich, UK) was used at 0.3 mM final concentration. Phosphate-buffered saline (PBS; Sigma-Aldrich, UK) was used to wash and dilute bacterial cells. Minimal media were prepared from 5x M9 salts (Sigma-Aldrich, UK), supplemented with 1 mM MgSO_4_ (Sigma-Aldrich, UK) and either 0.2 % (v/v) glycerol (Sigma-Aldrich, UK) or 0.2 % (w/v) casamino acids (Fisher Scientific, UK).

### DNA manipulation and cloning

Plasmids (**Supplementary Table 2**) were purified from overnight bacterial cultures using the Monarch Plasmid Miniprep Kit (New England Biolabs, UK). Polymerase chain reaction was carried out using Q5 high-fidelity polymerase (New England Biolabs, UK) for cloning or AppTaq RedMix (Appleton Woods, UK) for colony screening. Oligonucleotide primers (**Supplementary Table 3**) with and without modifications were purchased from Eurofins Genomics (Germany). Amplicons were purified using the Wizard SV Gel and PCR Clean-Up System (Promega, UK). Genomic DNA was purified using the Wizard® Genomic DNA Purification Kit (Promega, UK). Plasmids were assembled using NEBuilder HiFi DNA Assembly (New England Biolabs, UK).

### Conjugative mutagenesis

Deletion of the *ccdAB* and *aggR* genes from pAA_1070_ was achieved through conjugative mutagenesis since other methods (e.g. lambda Red recombineering) failed to produce satisfactory results in *E. coli* 1070/13. Briefly, isothermal DNA assembly was used to produce variants of pCONJ5K [29] containing a chloramphenicol resistance cassette flanked by pAA DNA sequences surrounding either *ccdAB* or *aggR*. *E. coli* MFD*pir* Δ*hsdR* carrying either plasmid variant was used as a donor strain for conjugation into recipient *E. coli* 1070/13. Donor strains were grown overnight in LB + DAP + chloramphenicol. The recipient strain was grown overnight in LB. From each, 1 ml culture was pelleted (16,000 x *g* for 2 min) and washed with an equal volume of PBS. 20 μl donor was mixed with 20 μl recipient and the total volume was spotted onto L-agar + DAP. When dry, the plate was incubated at 37 °C with aeration for four hours. Following incubation, bacteria were resuspended in 1 ml LB and one tenth of the volume was plated onto L-agar containing chloramphenicol for incubation overnight at 37 °C. Colony PCR was performed to check for the first crossover event and positive colonies were grown overnight in LB + chloramphenicol. From this, 1 ml culture was pelleted and washed three times in LB, then a further 4 ml LB was added and the culture was grown for 4 hours at 37 °C with aeration. From the incubated culture, 5 μl was plated onto sucrose agar and incubated overnight to obtain second crossover mutants (i.e. plasmid backbone excision). Final mutants were screened by colony PCR and Sanger sequencing.

### DNA sequencing and analysis

PCR amplicon sequencing for verification of clones and mutants was performed by Source BioScience (UK). Whole-genome hybrid Illumina/Nanopore sequencing of strains 1070/13 and 1070/13 pAA^−^ was performed by MicrobesNG (UK). MicrobesNG prepared Illumina sequencing libraries using the Nextera XT Library Prep kit (Illumina, UK) prior to sequencing on an Illumina NovaSeq 6000, for 2×250 bp paired-end reads. Adapters were trimmed from reads using Trimmomatic v0.30 with a sliding window quality cut-off of Q15 [30] and a minimum read length of 36 bp. Nanopore sequencing libraries were prepared using the Oxford Nanopore SQK-LSK109 kit with Native Barcoding kits EXP-NBD104/114, prior to loading on a GridION with a FLO-MIN106 (R.9.4.1) flow cell. Raw data was basecalled using the model 2021-05-17_dna_r9.4.1_minion_384_d37a2ab9.

Hybrid genomes were assembled in-house according to Eladawy *et al .*[31]. Briefly, end and middle adapters were trimmed from Nanopore sequencing reads using Porechop v0.2.4 (https://github.com/rrwick/Porechop) with thresholds of 95% and 85% respectively. Filtlong v0.2.1 was used to remove short reads, <1 kbp. Overlapping reads were assembled into complete circular molecules using Flye v2.9.2 [32], prior to being polished with both Nanopore and Illumina reads. Assembled contigs were first polished with Nanopore reads, using four iterations of Racon v1.5.0 [33] and one iteration of Medaka v1.8.0. Subsequently, Illumina reads were used to polish the resulting sequences, using Polypolish v0.5.0 [34], POLCA from the MaSuRCA v4.1.0 package [35] and Nextpolish v1.4.1 [36].

### Toxicity assay

To test the effect of toxins and antitoxins cloned onto pBAD33 and pGM101_neo_ derivatives respectively, the following protocol was used as previously described [26]. Strains carrying test plasmids or empty vector controls were grown overnight in LB + antibiotics + 0.2 % (w/v) glucose (Sigma-Aldrich, UK) and then subcultured to a starting optical density at 600 nm (OD_600nm_) of 0.01 in fresh media. Upon re-growing to OD_600nm_ = 0.1 – 0.2, bacterial cells were pelleted by centrifugation at 4,000 x *g* for 10 minutes and the supernatant discarded. Cells were resuspended in pre-warmed LB + antibiotics + 1 % (w/v) arabinose (Sigma-Aldrich, UK). Immediately and at timed intervals thereafter, cultures were serially diluted and plated onto L-agar containing antibiotics + 0.2 % (w/v) glucose, then incubated at 37 °C overnight to observe bacterial viability.

### Human cell culture and adhesion assay

HT29 cells were grown in Dulbecco’s modified eagle’s medium with high glucose (DMEM) (Sigma-Aldrich, UK), 10 % (v/v) fetal bovine serum (FBS) (Fisher Scientific, UK), 2 % (v/v) HEPES (v/v) (Sigma-Aldrich, UK) and 10 μg/ml PenStrep (Fisher Scientific, UK) in a T75 flask. Caco-2 cells were grown in DMEM and 10 % (v/v) FBS. Both cell lines were grown in a humidified incubator with 5 % CO_2_ at 37 °C. Cells were harvested after reaching approximately 80 % confluency by adding 2 ml TrypLE Express (Fisher Scientific, UK). Cells were seeded at 10^5^ cells per well in a 24 well plate the day before infection. All work on HT29 cells was carried out at the University of Oxford; experiments using Caco-2 cells were carried out at Nottingham Trent University.

For the adhesion assay, bacteria were grown overnight then subcultured in 10 ml LB to OD_600nm_ = 0.01 and re-grown to OD_600nm_ = 0.6. Cultures were then centrifuged at 4,000 x *g* for 5 min and the pellet resuspended in PBS before a further 2 min centrifugation. The pellet was then resuspended in pre-warmed DMEM to the volume harvested (taking into account culture removed for OD measurements). From this, the starting inoculum was serially diluted and plated onto L-agar for confirmation. The 24 well plate containing a confluent monolayer of eukaryotic cells was checked via microscopy before the growth media was removed from each well. Wells were then inoculated with 1 ml DMEM containing 10^7^ bacterial cells per well (MOI = 100) in triplicate. An additional well per strain that was not seeded with eukaryotic cells was inoculated with bacteria to account for any adherence of bacteria to the surface of the well itself, the result of which was subtracted from test wells. The 24-well plate containing the infected eukaryotic cells was then incubated at 37 °C, 5% CO_2_ for either 30 minutes or 3 hours. Following incubation, media was removed from all wells and wells were washed thrice with 1 ml PBS to remove any non-adhered bacteria. 1 ml DMEM containing 1 % (v/v) triton X-100 (Promega, UK) was then added to each well with vigorous pipetting to detach and disrupt cells from the well surface. Samples from each well were then diluted and plated on L-agar and incubated at 37 °C overnight. Results were normalised to the starting inoculum.

### Biofilm assay

Overnight cultures were diluted to OD_600nm_ = 0.5 in PBS. 1 ml growth medium was added to each well in a 24-well plate to which 100 μl diluted culture was inoculated in triplicate. Control (blank) wells contained growth medium only. The 24-well plate was incubated at 37 °C for 18 hours. Growth medium was carefully removed by tipping and blotting. Wells were then washed with 1 ml PBS by shaking at 100 rpm for 5 minutes. PBS was then removed and the wash repeated. Each well was stained with 500 μl of 0.1 % (w/v) crystal violet dye (Sigma-Aldrich, UK) and was incubated at room temperature for 1 hour. Excess dye was then drained, and wells were washed gently with 1 ml PBS and then tipped and blotted. This wash step was then repeated until the PBS remained colourless when added to the wells. To solubilise the dye, 200 μl 70% (v/v) ethanol (Fisher Scientific, UK) was added to each well and the plate was shaken for 2 minutes at 100 rpm. Subsequently, 100 μl was taken from each well into a 96-well plate and the OD measured at 540 nm. Values from the blank wells were subtracted to account for well-only staining.

### Motility assay

To assess motility, a single colony was picked via sterile needle from a streak grown overnight, then used to stab the centre of LB + 0.3 % (w/v) agar containing 1 % (w/v) triphenyltetrazolium chloride (Sigma-Aldrich, UK). Plates were incubated at 37 °C for 18 hours, then motility diameter around the stab point was measured.

### Digital droplet PCR (ddPCR)

For ddPCR, the phenol-chloroform method of DNA purification was used. 600 μl overnight bacterial culture was added to a phase lock tube. One volume of phenol:chloroform:isoamyl alcohol (25:24:1) (Sigma-Aldrich, UK) was added to the sample and vortexed. The sample was centrifuged at room temperature for 5 min at 16,000 × *g*. The upper aqueous phase was transferred to a fresh tube. Reagents were added to the aqueous phase in the following order: 1 μl glycogen 20 μg/μl (Sigma-Aldrich, UK); 0.5x sample volume 7.5 M ammonium acetate (Sigma-Aldrich, UK); and 2.5x sample volume 100% ethanol. DNA precipitated overnight at -20 °C. The sample was then centrifuged at 4 °C for 30 min at 16,000 × *g.* The supernatant was removed and 150 μl of 70 % (v/v) ethanol added. The sample was centrifuged at 4 °C for 2 min at 16,000 × *g* and ethanol removed. The DNA pellet was dried at room temperature for 10 min before being resuspended in 30 μl of nuclease-free water and concentration measured spectrophotometrically.

The QX200 Droplet Digital PCR System and recommended consumables (Bio-Rad, UK) were used for all ddPCR experiments. Primers and probes were designed using Primer3plus [37] with probes modified at 5’ end with fluorophore FAM for the reference gene and HEX for the plasmid marker and modified at the 3’ end with the quencher BHQ-1 (Eurofins Genomics, Germany).

The master mix was made as described in **Supplementary Table 4** and 20 μl added to each well in the DG8 cartridge, followed by 70 μl QX200 droplet generation oil for probes. The cassette was placed into the DG8 Cartridge holder and DG8 Gasket added. Droplets were then formed using the QX200 Droplet Generator. The resulting droplets were carefully transferred to a ddPCR 96-well plate and PCR plate pierceable heat seal foil added and sealed using the PX1 PCR plate sealer. PCR was then carried out using the C1000 Touch thermocycler with 96-deep well reaction module with the following PCR conditions: enzyme activation 95 °C for 10 minutes; 40 cycles of denaturation at 94 °C for 30 seconds and annealing/extension 55-64 °C for 1 minute; 1 cycle of enzyme deactivation at 98 °C for 10 minutes. Results were analysed using QuantaSoft software version 1.7.4.0917.

### Statistical analysis

All statistical analysis was carried out in GraphPad Prism [38]. Data were log-transformed prior to analysis by either one- or two-way ANOVA or t-test depending upon the experimental requirements. Where relevant, appropriate post-tests were used to interrogate data further. Individual tests for each experiment are detailed in the figure legends. For plasmid loss assays, statistically significant outliers were identified and removed via Grubbs test; doing so did not change the overall conclusion of the experiments.

## RESULTS

## pAA has a low plasmid copy number typical of other *E. coli* virulence plasmids

*E. coli* O104:H4 strain 1070/13 is representative of the O104:H4 isolates recovered from a multi-pathogen Newcastle outbreak [28]. Strain 1070/13 underwent Illumina whole-genome sequencing after its isolation by Dallman *et al*. and was found to contain a variant of pAA, hereafter referred to as pAA_1070_. To confirm the plasmid sequence, we carried out hybrid genome sequencing of the isolate and revealed an intact plasmid (accession number CP171475) that is 99.95% identical across 88% of the plasmid sequence to pTY2 from the 2011 outbreak strain TY2482 [4,5]. Differences between pAA_1070_ and pTY2 primarily consist of insertion sequence elements such as ISKpn26-, ISEc43- and IS66-related ORFs in pAA_1070_ that appear as assembly gaps in the pTY2 sequence. To obtain an approximate plasmid copy number for pAA_1070_, we analysed Dallman and colleagues’ short-read sequence coverage of strain 1070/13 separately mapped to both the pTY2 plasmid reference sequence [5] and the *E. coli* K-12 MG1655 chromosome sequence [39]. The former gave a mean read depth of 264.6x, and the latter a mean read depth of 100.3x, indicating a plasmid copy number of approximately 2.6 plasmid molecules per chromosome; a result expected from prior studies on enteric virulence plasmids [40–43]. Long read sequencing from our hybrid approach agreed, giving a copy number of 2.48 plasmids per chromosome (calculated from an overall plasmid contig depth of 2.82x to a mean chromosome contig depth of 1.138x).

To confirm this value experimentally, we carried out droplet digital polymerase chain reaction (ddPCR) to determine the absolute DNA quantities of a range of plasmid genes relative to chromosome genes in *E. coli* 1070/13 grown to stationary phase in LB at 37°C. Results (**Figure 1A**) show an estimated plasmid copy number of between approximately 1.5 and 3.3 depending upon probe combination; the mean of 2.2 plasmid molecules per chromosome was in general agreement with the copy number extrapolated from sequencing data. As a control for the ddPCR assay, we also produced a pAA-cured variant of strain 1070/13. Contrasting the apparent instability of EAEC plasmids reported *in vivo* [44], pAA_1070_ *in vitro* was remarkably stable relative to the virulence plasmids of organisms such as *Shigella* [26] and curing required multiple heat-shock treatments. The resulting strain was designated 1070/13 pAA^−^ and was negative for plasmid probe fluorescence in ddPCR assays, whereas the wild type remained positive (**Figure 1B-C**).

**Figure 1:**
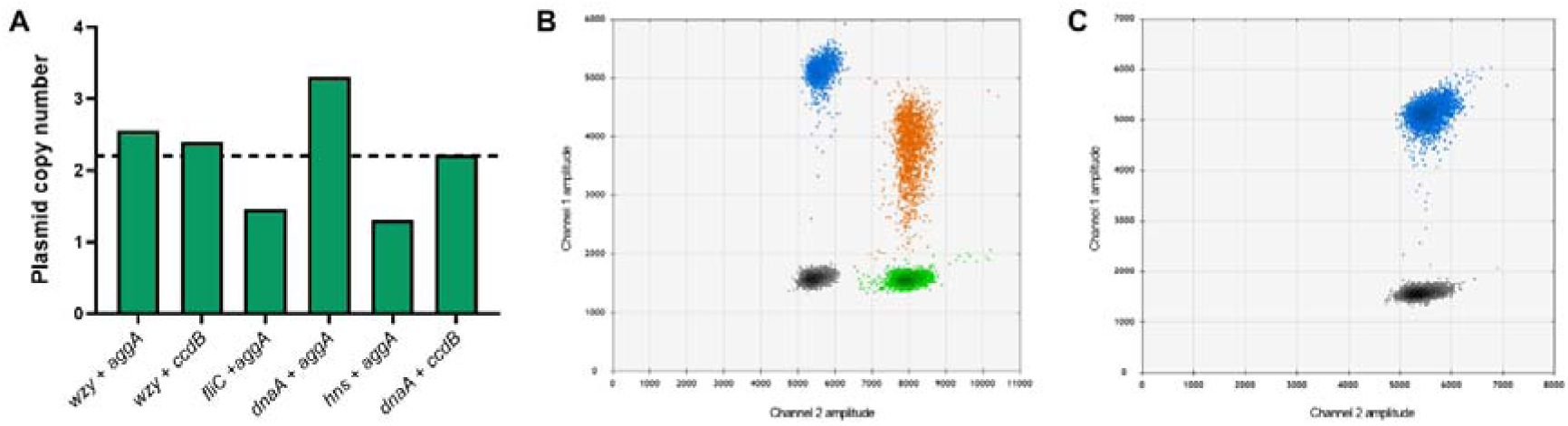
Determination of pAA_1070_ copy number by ddPCR. **(A)** Plasmid copy number per chromosome, calculated from different probe combinations as indicated. Experiments performed at annealing temperature 61.2 °C with DNA template concentration 50 ng/μl. Probes targeting chromosomal genes (*wzy, fliC, dnaA* and *hns*) labelled with FAM. Probes targeting plasmid genes (*aggA* and *ccd B*) labelled with HEX. No template controls showed double-negative results for all probe combinations (not quantified on graph). Dotted line: mean pAA_1070_ copy number calculated from all probe combinations. **(B, C)** Representative 2D amplitude plots of ddPCR assay using *wzy* and *aggA* probes and template DNA from either **(B)** 1070/13 or **(C)** 1070/13 pAA^−^. Grey dots: double-negative droplets; blue dots: FAM-positive droplets (*wzy* only); green dots: HEX-positive droplets (*aggA* only); orange dots: double-positive droplets (*wzy* and *aggA*).

### pAA carries at least one functional toxin-antitoxin system

Bioinformatic analysis of pAA_1070_ revealed the presence of multiple putative TA genes. Immediately adjacent to the FII origin of replication are two operons putatively encoding RelE/ParE-like toxins and their cognate antitoxins. In addition to this, a second TA-related locus approximately 5-7 kb from the origin contains two divergently-transcribed TA operons putatively encoding CcdAB and VapBC. Interestingly, in addition to the expected ATG start codons, BASys annotation [45] predicted alternative upstream start codons for both toxin genes (*ccdB*: TTG and *vapC*: GTG). We noted that VapC includes a specific active site residue (N107, counted from the GTG start codon) that may render the toxin nonfunctional [46].

In order to analyse the roles of these systems, we first cloned the individual toxin genes into an arabinose-inducible expression vector, pBAD33 [47], and assessed the resulting protein products for toxicity in *E. coli* DH5α. For *ccdB* and *vapC*, since both of the alternative start codons seem to be associated with a putative Shine-Dalgarno sequence and alternative start codons are linked to translation under stress [48–50], we included both variants of each gene in our initial toxicity screen, using the nomenclature CcdB_101_/CcdB_110_ and VapC_133_/VapC_142_ referring to the toxin length in amino acids. We surmised that, regardless of whether or not the alternative start codons were functional, the longer-length constructs would give the best chance of observing toxicity (barring post-transcriptional control), whereas the shorter-length constructs could reveal any requirement for an extended N-terminal sequence.

Results (**Figure 2A**) showed no significant toxicity 180 minutes post induction of either VapC or the two RelE/ParE-like toxins (*P* ≥ 0.285 compared to the empty vector control), whereas both CcdB variants were significantly toxic (*P* < 0.0001). Interestingly, there was also a minor but significant difference between the effect of the two CcdB variants when compared to one another at 30 minutes (*P* = 0.0024) and 180 minutes (*P* = 0.0178) post induction; though, as this effect did not persist between *E. coli* strains (see below) and we hence did not verify the proteins produced by our constructs, we do not intend to suggest here that this result is due to the production of an alternate protein variant *per se*.

**Figure 2:**
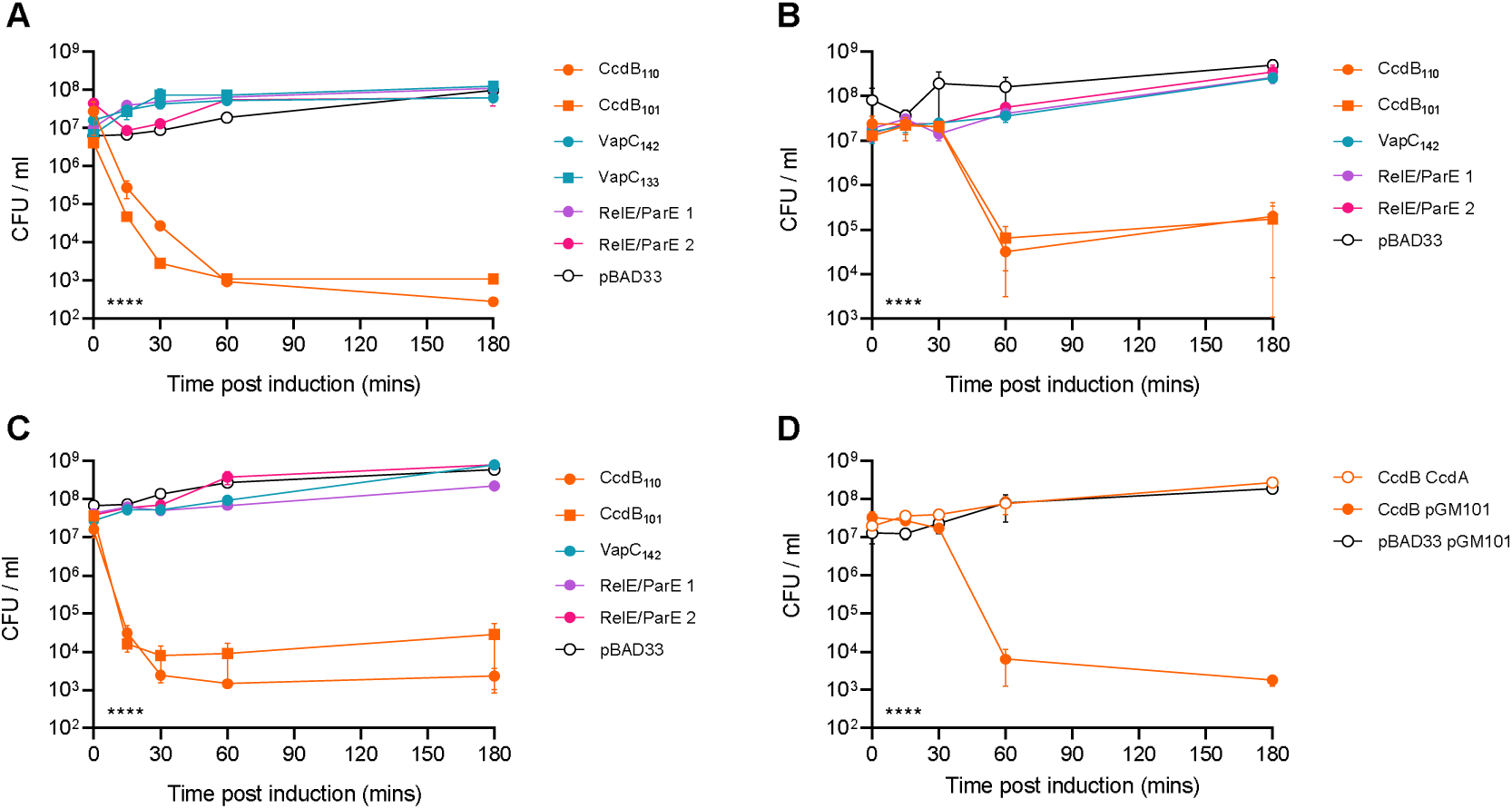
Characterisation of TA systems on pAA _1070_. Mean bacterial viability after induction of toxin production from pBAD33; **(A)** in *E. coli* DH5α grown in LB; **(B)** in *E. coli* 1070/13 pAA^−^ grown in LB; **(C)** in *E. coli* 1070/13 pAA^−^ grown in M9 with casamino acids; **(D)** in *E. coli* 1070/13 pAA^−^ grown in LB in the presence/absence of cognate antitoxin produced from pGM101_neo_. Empty vector controls in each case are indicated as pBAD33 and/or pGM101. Error bars show standard error of the mean. Statistical analysis by two-way ANOVA with Tukey’s multiple comparison tests: indicator in lower left shows the overall effect of strain in each experiment; **** *P* < 0.0001 (from *n* = 3 biological replicates).

To confirm toxicity in the native host strain in the absence of any antitoxin interference from the native pAA plasmid, 1070/13 pAA^−^ was used in a toxicity assay under the same experimental conditions as *E. c o l*D*i*H5α (**Figure 2B**). Due to the potential difference between putative CcdB variants seen in DH5α, we included both CcdB_101_ and CcdB_110_ in this experiment, whereas we included only the longer-length VapC_142_ clone. Production of either CcdB variant again significantly reduced cell viability, with the most dramatic effect at 60 minutes post induction (*P* < 0.02 relative to the empty vector control). In this strain background, there was no significant difference between the two CcdB variants (*P* ≥ 0.96 throughout the assay). As in DH5α, there was no significant toxicity observed when expressing any of the non-*ccdB* genes (*P* > 0.57).

Suspecting that the effect of the TA systems might be impacted by growth media, we also tested the toxicity of the above constructs during growth in M9 minimal media supplemented with casamino acids (**Figure 2C**). Results were similar to those observed during growth in LB, with a significant effect of both CcdB variants on viability (most pronounced at 60 minutes post induction, *P* < 0.03). We also noted that both VapC and one of the RelE/ParE variants showed significant toxicity at this timepoint (*P* < 0.045) but the effect was very minor (a decrease in viability of less than 10-fold for each compared to more than 1,000-fold for CcdB) so this was not pursued further. As in LB, there was no significant difference between the CcdB variants throughout the assay (*P* > 0.39).

To verify that CcdAB functions as a cognate TA pair on pAA_1070_, the *ccdA* gene was cloned onto a second vector, pGM101_neo_, under control of its native promoter to allow for induction of gene expression via conditional cooperativity [51]. pGM101_neo_ was constructed as a neomycin-resistant derivative of the promoterless, pBAD33-compatible vector pGM101 [26], used to ensure selection in the ampicillin-resistant 1070/13 strain. A toxicity assay was then conducted in the 1070/13 pAA^−^ host background using CcdB_110_ as the representative toxin construct. As expected, expression of *ccdB* from pBAD33 in the presence of the *ccdA*-containing vector no longer resulted in loss of bacterial cell viability (**Figure 2D** ; *P* > 0.23 vs. empty vector control throughout), implying that the *ccdAB* operon on pAA_1070_ encodes a classical TA system.

Lastly, to investigate the mode of action of CcdB from pAA_1070_, we tested the toxicity of our pBAD33 constructs in two commercial CcdB-resistant strains: *ccdB* Survival 2 T1^R^ (Invitrogen; resistance genotype not known) and DB3.1 (originally from Invitrogen but no longer available; known to harbour the *gyrA462* allele). The *Shigella flexneri* M90T pINV variant of CcdB [26] was included as a control. Results showed that whilst all toxin variants reduced the viability of *ccdB* Survival cells relative to the empty vector (**Supplementary Figure 1A**, *P* ≤ 0.0012 at 180 minutes post induction), there was no toxicity in DB3.1 (**Supplementary Figure 1B**, *P* > 0.14 at 180 minutes post induction). This suggests that CcdB from pAA_1070_ targets DNA gyrase, as expected.

In order to assess the impact of the CcdAB module on plasmid maintenance, we replaced the *ccdB* gene with a chloramphenicol resistance cassette, rendering the TA locus non-functional. Counterselectable markers (e.g. *sacB*) have been used in TA research to successfully quantify very small levels of plasmid loss [26], but we were unable to generate such a pAA mutant, for reasons unknown. We therefore relied initially upon quantification of plasmid loss via PCR and/or plating to agar with and without chloramphenicol. Our experiments revealed no measurable destabilisation of pAA within the detection limit of our experiments, regardless of the presence or absence of *ccdAB* (<1% plasmid loss for both the WT and Δ*ccdB* strain).

Given the low detection limit of the above experiments to quantify plasmid loss, we constructed a stability test vector (“pSTAB”) with and without the *ccdAB* locus from pAA_1070_. pSTAB vectors contain a selectable neomycin resistance marker as well as a counterselectable *sacB* gene [52], improving the detection limit in experiments to approximately 0.0001% plasmid loss [53,54]. These vectors rely upon the native virulence plasmid replicon for replication and copy number control; interestingly, we were only able to obtain successful pSTAB clone (designated pMW_O104) by including the entirety of the 6,530 bp dual FII/FIB replicon from pAA_1070_, unlike a previous *Shigella* pINV study in which the 2,600 bp FII replicon alone was sufficient [54]. Quantifying pMW_O104 and pMW_O104::*ccdAB* plasmid loss over approximately 20-25 generations of growth, we determined that the CcdAB system provides a minor but statistically significant improvement in plasmid stability (**Figure 3**; *P* < 0.0001).

**Figure 3:**
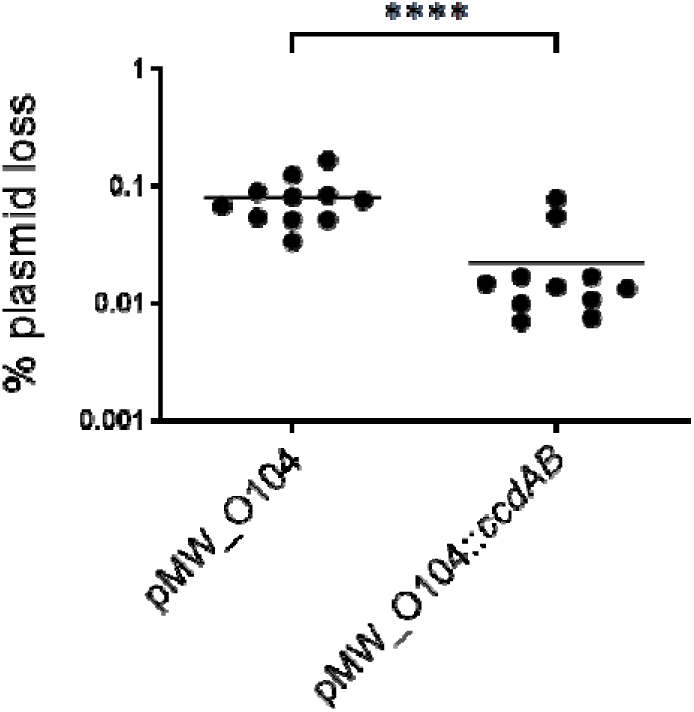
Effect of CcdAB on a pSTAB test vector containing the natural FII/FIB dual replicon region of pAA_1070_. Plasmid stability in E. coli NEB 10-beta determined by plating onto neomycin (plasmid-positive) or sucrose (plasmid-negative) in accordance with previous work [54,26]. Each point shows plasmid loss as a percentage of the total viable cell population within a single biological replicate (one independent colony grown on L-agar for 20-25 generations at 37 °C). Solid lines show the mean. Data combined from multiple independent experiments. Statistical analysis by t-test: **** P < 0.0001.

### pAA_1070_ influences adhesion to host cells but not biofilm formation

EAEC plasmid research has typically been carried out in TY2482 or other strains from the 2011 outbreak, which contain a Shigatoxigenic phage [3,6,44], or in strains with different variants of the plasmid, e.g. prototypical EAEC strain 042 [55–58]. In order to elucidate the phenotypic role of pAA in a pathogenic strain closely related to TY2482 but lacking the Shiga toxin, 1070/13 pAA^−^ and its parent strain were subjected to a range of tests to assess any change in growth rate, biofilm formation and adherence to intestinal cells *in vitro*.

Loss of pAA_1070_ resulted in no significant change in growth rate or growth yield for bacteria grown in either rich medium (LB), minimal medium supplemented with glycerol or minimal medium supplemented with casamino acids (**Supplementary Figure 2**). Similarly, there was poor biofilm production (**Figure 4A**) from the strain regardless of whether it contained the plasmid (pAA^+^ vs. pAA^−^, *P* = 0.9753) and O104:H4 biofilm production was similar to that of *E. coli* K-12 laboratory strain MG1655 (all O104:H4 strain derivatives vs. MG1655, *P* > 0.91), whereas a *Pseudomonas aeruginosa* PAO1 control [59] showed significantly higher biofilm production (PAO1 vs. all *E. coli* strains*, P* < 0.01).

**Figure 4:**
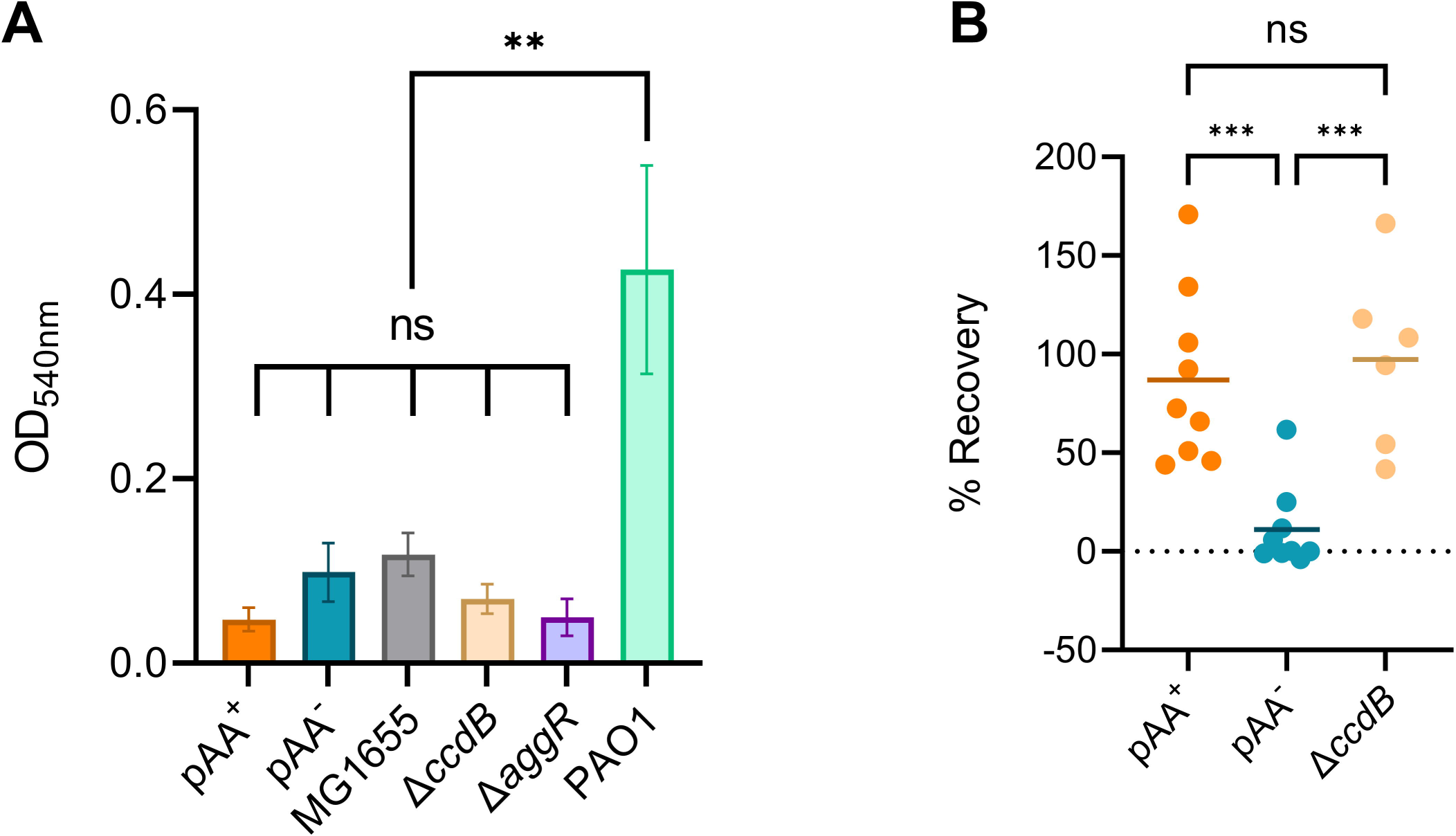
Effect of pAA on virulence-related phenotypes of O104:H4 strain 1070/13. **(A)** Mean biofilm production after 24 hours’ growth in LB at 37 °C, assayed via crystal violet staining. Control: *P. aeruginosa* PAO1. Error bars show standard error of the mean. **(B)** Per cent recovery of bacterial cells (normalised to inoculum) after incubation for 180 minutes with Caco-2 cells. Each point is one biological replicate. Solid lines show the mean. Statistical analysis by one-way ANOVA with Tukey’s multiple comparison tests: ns, not significant (P ≥ 0.05); ** *P* < 0.01; *** *P* < 0.001 (from n ≥ 3 biological replicates).

We next assessed the ability of 1070/13 and 1070/13 pAA^−^ to adhere to human colon epithelial cells *in vitro* . As expected, the loss of pAA_1070_ significantly reduced the organism’s ability to adhere to both Caco-2 cells (**Figure 4B** ; *P* = 0.0008) and HT-29 cells (**Supplementary Figure 3A**; *P* = 0.0067), most likely due to lack of AAF. This is consistent with previously-published work on other EAEC strains [60–62,58], so we did not investigate this particular phenotype further. However, we noted that the human cells co-cultured with wild type EAEC varied dramatically in their morphology from those co-cultured with the plasmid-free variant or the non-infected control (**Supplementary Figure 3B-D**). The Δ*ccdB* mutation did not affect biofilm formation (**Figure 4A**, *P* = 0.9995) or host cell adhesion (**Figure 4B**, *P* = 0.8585) relative to the wild type strain.

### Motility in strain 1070/13 can be abolished by excision of a chromosomal insertion sequence

In order to assess the impact of pAA on another putative virulence phenotype, we compared the motility of 1070/13 pAA^−^ with that of the wild type strain. The plasmid-free strain was significantly less motile than the wild type at both human host (37 °C) and non-host (30 °C) growth temperatures (**Figure 5A**, *P* < 0.0001). To explore two possible reasons for this, we tested the motility of our Δ*ccdB* mutant as well as a strain lacking the plasmid-encoded regulator AggR. Whilst we observed no motility change in the Δ*aggR* strain (*P* > 0.2 for both temperatures), to our surprise, our Δ*ccdB* mutant showed the same lack of motility as the plasmid-free strain (Δ*ccdB* vs. pAA^+^, *P* < 0.0001). Wild type O104:H4 proved significantly more motile than MG1655 regardless of temperature (*P* < 0.0001).

**Figure 5:**
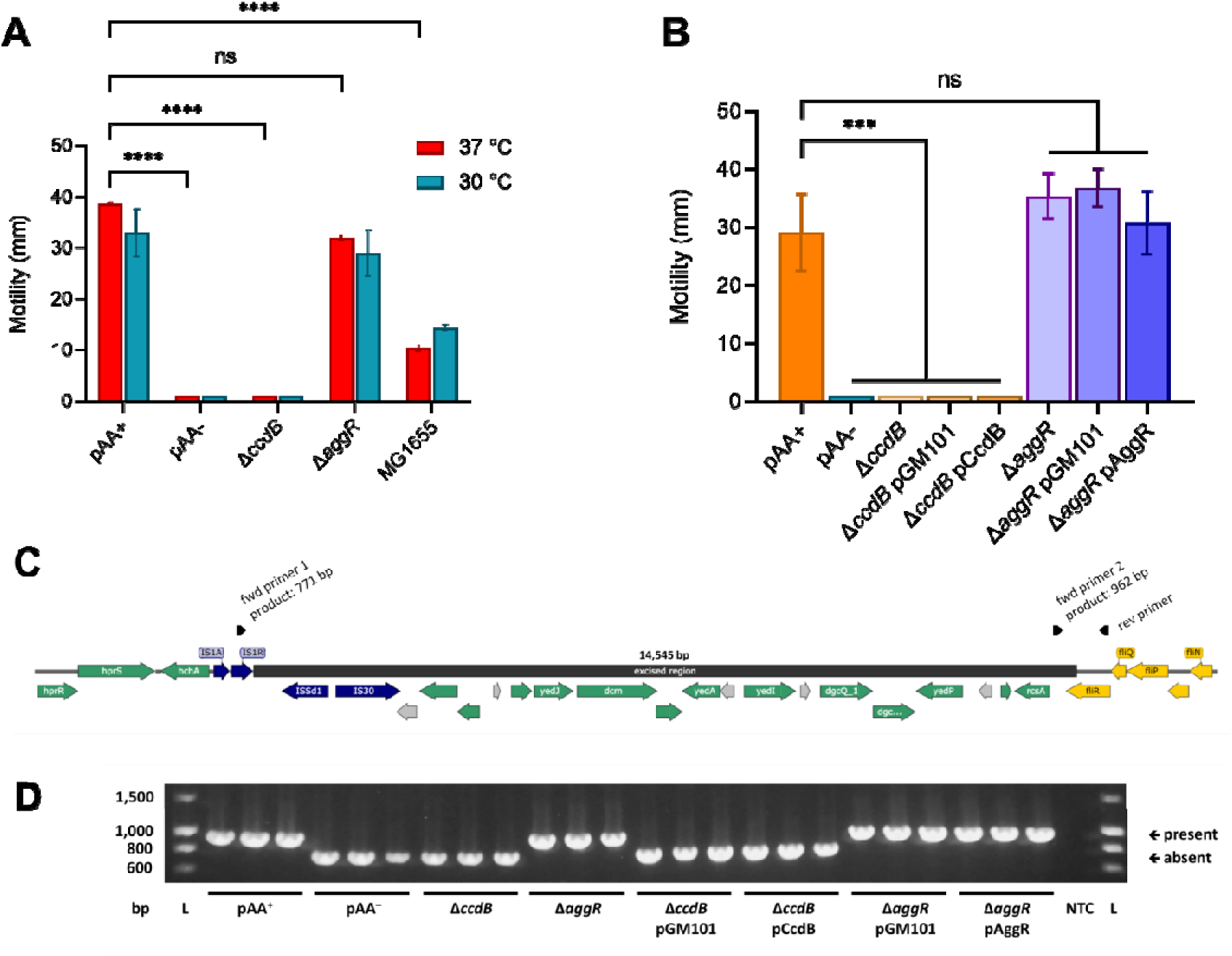
Effect of pAA on motility of O104:H4 strain 1070/13. **(A)** Mean distance from stab point in motility agar at 37 °C (red) and 30 °C (blue). **(B)** Mean distance from stab point in motility agar at 37 °C with and without the empty vector control (pGM101) or complementation plasmids (pCcdB or pAggR). For both experiments, a lack of evident motility was entered as 1 mm for visualisation/analysis purposes. Error bars show standard error of the mean. Statistical analysis by one- or two-way ANOVA with Tukey’s or Sidak’s multiple comparison tests as appropriate: ns, not significant (P ≥ 0.05); *** P < 0.001; **** P < 0.0001 (from n = 3 biological replicates). **(C)** Schematic of 14,545 bp region putatively excised from the 1070/13 chromosome, bordering fliR. Dark blue: transposase-related genes; yellow: flagellar genes; green: other genes of known function; grey: genes encoding hypothetical proteins. PCR primers indicated by black arrows; rev primer forms a product of either 771 bp (pairing with fwd primer 1, indicating excision of region) or 962 bp (pairing with fwd primer 2, indicating presence of region). **(D)** PCR screen of bacteria used in motility assay (three replicates per strain), indicating presence/absence of the excised region. Representative of multiple independent PCR/motility assays. NTC: no template control. L: ladder; band sizes (bp) shown left.

Supposing that the CcdAB complex might have a regulatory function that had been disrupted in our mutant, we constructed a complementation plasmid consisting of pGM101_neo_ carrying the whole *<u>ccdAB</u>* operon plus its promoter, and tested motility at 37 °C in strains carrying either this construct or the empty vector (**Figure 5B**). The complementation construct was unable to restore motility to the mutant strain (pCcdB vs. pGM101_neo_ in Δ*ccdB* background, *P* > 0.9999), suggesting either that the construct somehow failed to compensate for the deletion despite it being a genuine effect, or that lack of motility in the Δ*ccdB* mutant is due to an off-target mutation. A similar complementation plasmid carrying *aggR* had no impact on motility (pAggR vs. pGM101_neo_ in Δ*aggR* background, *P* = 0.9241).

To confirm the reason for lack of motility, we analysed the genome sequence of 1070/13 and 1070/13 pAA^−^ strains. This revealed the absence of a chromosomal region bordering IS1, IS3 and IS30 transposase genes (**Figure 5C**) in the plasmid-free strain relative to the parent isolate, resulting in substitution of the C-terminal 64 amino acids (∼25%) of flagellar export apparatus component FliR with a nonhomologous sequence of 27 amino acids. Based on this information, we designed primers (**Figure 5C**) to detect the configuration of this genomic region and confirmed that motility loss in all instances, including the Δ*ccdB* mutant, correlated perfectly with excision of this region, whereas strains containing the sequence (and hence the full-length *fliR* gene), including the Δ*aggR* mutant, remained motile. We also noted that the wild type isolate sometimes spontaneously became non-motile during laboratory culture and that this had indeed occurred during growth of the isolate for hybrid genome sequencing – hence, the sequence we provide herein for the 1070/13 parent strain differs from the Illumina data produced by Dallman *et al.* [28] in the absence of the above genomic region. However, in all instances where the strain remained motile throughout a given experiment, PCR confirmed the region’s presence. Representative PCR data of all strains is shown in **Figure 5D**. Importantly, we were also able to confirm from our prior assays that this motility defect affects neither biofilm formation nor host cell adhesion (**Figure 4**; Δ*ccdB* mutant).

## Discussion

We have shown in this study that the CcdAB TA system on pAA_1070_ is a functional toxin and antitoxin pair, with CcdB likely targeting DNA gyrase, but has only a minor effect on plasmid stability under the conditions tested. We have also shown almost no measurable difference in toxicity between constructs encoding CcdB_101_, the traditionally-characterised variant [63,64], and CcdB_110_ (a variant putatively translated from an alternative TTG start codon on pAA_1070_). However, since we saw no difference in toxicity against the native host bacterium, we did not experimentally validate the proteins produced from each construct. It therefore remains possible (even likely) that the TTG start codon is not genuinely recognised by the translation machinery, that it is an annotation error, and that our CcdB_110_ construct is simply producing the CcdB_101_ variant. If ongoing work suggests a potential impact of this alternate start codon, further protein characterisation should be performed as a priority.

Divergently-transcribed TA systems are not well-studied, and given the role of conditional cooperativity, i.e. binding of type II TA complexes to their own promoters, we wondered whether the CcdAB and VapBC systems on pAA_1070_ might impact one another’s gene expression as has been reported for several TA systems [65–67] and the divergently-transcribed *par1*/*par2* partitioning loci on pB171 [68]. However, with the finding that this particular VapC variant is non-toxic and that this is likely due to an active site mutation [46], this avenue was not pursued further in this particular work. Nonetheless, regardless of a lack of VapC toxicity, putative regulation by the VapBC complex may also alter the effectiveness of the CcdAB system with regards to plasmid stability. Notably, the strain we used to perform pSTAB analysis does not encode VapBC. Though we did demonstrate a significant *ccdAB*-dependent increase in pSTAB_1070_ stability, the effect was not dramatic (< 10-fold). This subtle plasmid-stabilising role is not atypical of CcdAB TA systems, where some members of this particular TA system family have been shown to impart little or no stabilising effect overall under laboratory conditions despite being highly toxic [26,27]. It is possible that the pAA CcdAB system functions primarily not as a PSK module but as a regulator of other cellular functions. The regulatory function of both non-toxic and toxic but non-stabilising TA complexes is currently a subject of ongoing investigation within our laboratory.

We have shown that pAA is a low-copy virulence plasmid, similar to other pathovar-defining plasmids in *E. coli* such as pINV (copy number approximately 1 per chromosome [41]). We have also confirmed that pAA_1070_ is remarkably stable *in vitro*, with <1% natural plasmid loss occurring over multiple subcultures in liquid media, in agreement with a stress study on the 2011 outbreak strain of hybridised EAEC/EHEC [69] and consistent with our pSTAB results (approximately 0.1% plasmid loss over 20-25 generations of growth on solid medium). The lack of a stabilising TA system on this plasmid is therefore curious, as such maintenance elements are important for the stability of similar virulence plasmids [70]. We note that the plasmid carries genes related to ParA- and ParM-type partitioning systems; such systems function by physically segregating plasmid molecules during cell division [71] as opposed to the PSK activity of TA systems. TA systems and partitioning systems often work synergistically to optimise plasmid maintenance without excessive PSK [72]. However, the partitioning operons on pAA_1070_ are incomplete and are therefore unlikely to contribute to plasmid maintenance. Instead, as suggested by our pSTAB assay, the dual FII/FIB replicon of pAA_1070_ may be the cause of the plasmid’s *in vitro* stability. Indeed, it was not possible for us to generate pSTAB vectors using the FII replicon alone, contrasting past success with other IncFII and IncA/C plasmids [54,73]. We also note here the work by Zhang *et al*. [44] related to Stx-encoding O104:H4 in case patients, which showed that pAA may be far less stable *in vivo* than *in vitro*. It is possible that laboratory conditions are insufficient to reflect the host environment and that time-dependent pAA loss during infection is a common factor in human EAEC pathogenicity, though this would likely render the bacterium relatively non-pathogenic after passage through the host. Alternatively, pAA loss within the host may be correlated to production of Shiga toxin. To our knowledge, no research has addressed pAA loss in non-Shigatoxigenic EAEC isolates during the process of infection within patients.

Studies have previously reported the enhanced motility of EAEC strains compared to MG1655 [62], agreeing with our data. The plasmid-encoded EAEC virulence regulator AggR may negatively regulate the aggregation factor antigen 43 [74] and positively regulate flagellar genes [75]. In the former work, no motility effects were observed in the *aggR* mutant background, in agreement with our study. Contrasting this, motility was increased by overexpressing *aggR* in EAEC strain 042 [75]. This did not occur with the addition of our *aggR* complementation vector; though, as the *aggR* promoter is under complex regulation, increasing the gene’s copy number may not increase AggR production *per se*, or there may be relevant differences between EAEC strains. Notably, EAEC strain 042 carries a larger pAA variant than strain 1070/13 (113.3 kb vs 75.5 kb) and encodes AAF/II rather than AAF/I [55,28]. Indeed, Prieto *et al.* [75] showed that motility in strain 042 was increased by loss of the pAA plasmid, which we cannot rule out due to conflicting effects of *fliR* mutation in our pAA^−^ strain.

We demonstrated that loss of motility in EAEC strain 1070/13 can occur readily during laboratory culture and mutagenesis. This phenotype was found to correlate perfectly with the spontaneous excision of a chromosomal region bordered by transposase genes and insertion sequences, containing the 3’ end of the *fliR* gene. We note this as a cautionary tale in the era of readily-accessible whole genome sequencing; knowledge of the excised region in advance of our motility and complementation experiments would have been an advantage. The same chromosomal region, including IS elements, is present in the 2011 outbreak strain TY2482 (shotgun sequence scaffold4; accession NZ_GL989603.1) [5], though it is unknown if motility loss has ever occurred during experiments with that strain. Whilst reversible motility loss via transposon excision and re-insertion may reflect a type of phase variation in EAEC, we did not observe reversion to a motile phenotype during our assays. Alternatively, the ability of the organism to lose motility may confer an evolutionary benefit and future route to heightened pathogenesis, as some other enteric pathogens such as *Shigella* contain cryptic flagellar genes [76]. We observed no impact of this motility loss on either biofilm formation or host cell adhesion in our experiments. Ultimately, our study reflects the complexity of working with pathogenic outbreak isolates rather than laboratory strains, and validates the importance of doing so.

## Abbreviations

AAF: aggregative adherence fimbriae;
ddPCR: digital droplet polymerase chain reaction;
EAEC: enteroaggregative *Escherichia coli* ;
EHEC: enterohaemorrhagic *Escherichia coli*;
FAM: fluorescein;
HEX: hexachlorofluorescein;
MGE: mobile genetic element;
pAA: aggregative adherence plasmid;
pINV: invasion plasmid;
PSK: postsegregational killing

## Author Contributions

RW: Investigation, methodology, conceptualisation, data curation, visualisation, writing (review & editing). MC: Investigation, methodology, conceptualisation, data curation, writing (review & editing). BD: Supervision, resources, writing (review & editing). JCT: Formal analysis, methodology, writing (review & editing). GM: Investigation, methodology, conceptualisation, data curation, visualisation, writing (original draft), writing (review & editing), formal analysis, project administration, supervision, funding acquisition.

## Acknowledgements

We thank Claire Jenkins for strain provision under a Material Transfer Agreement (2017-03-23) from Public Health England (now UKHSA). We are grateful for the enthusiastic support of the Tang laboratory (University of Oxford) for essential training and strain provision. We also thank Jordan Shutt-McCabe for research assistance during the project, as well as members of AROM (NTU) for helpful discussions.

---

**Supplementary Figure 1:**
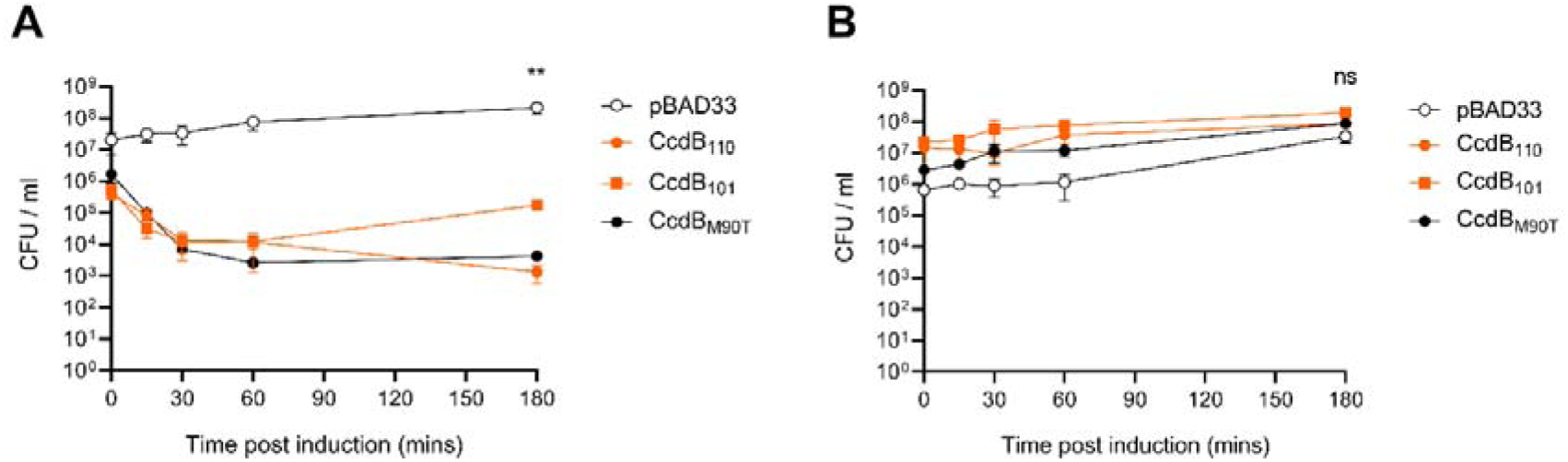
Toxicity of CcdB from pAA_1070_ in CcdB-resistant cloning strains. Mean bacterial viability after induction of toxin production from pBAD33; **(A)** in *E. coli ccdB* Survival 2 T1^R^; **(B)** in *E. coli* gyrase mutant DB3.1. Empty vector control in each experiment is indicated as pBAD33. Error bars show standard error of the mean. Statistical analysis by two-way ANOVA with Dunnett’s multiple comparison tests: ns, not significant (*P* > 0.05); ** *P* < 0.01 (from *n* = 3 biological replicates).

**Supplementary Figure 2:**
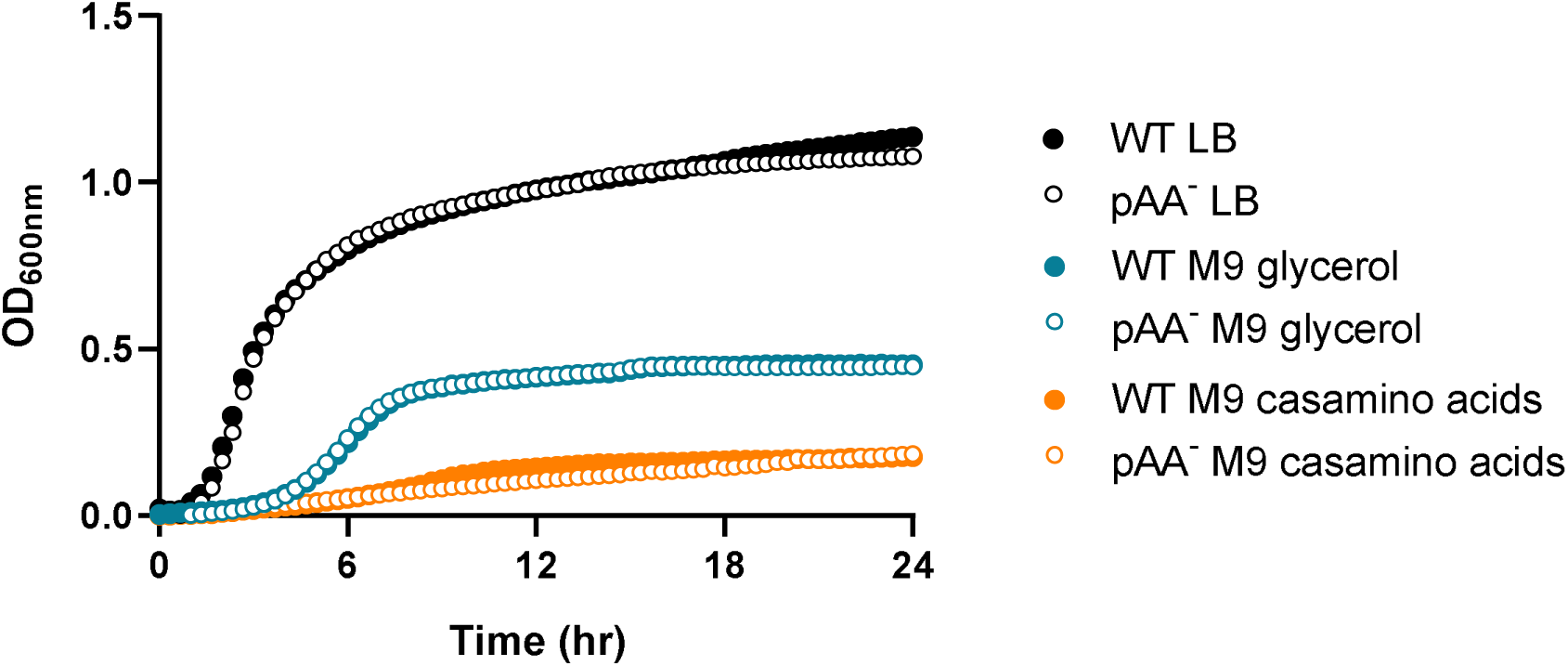
Growth of O104:H4 strain 1070/13 (WT) and its plasmid-free derivative (pAA^−^) in various laboratory media. OD _600nm_ measured every 20 minutes at 37 °C with aeration in 24-well plate. Mean of two biological replicates shown (from six technical replicates).

**Supplementary Figure 3:**
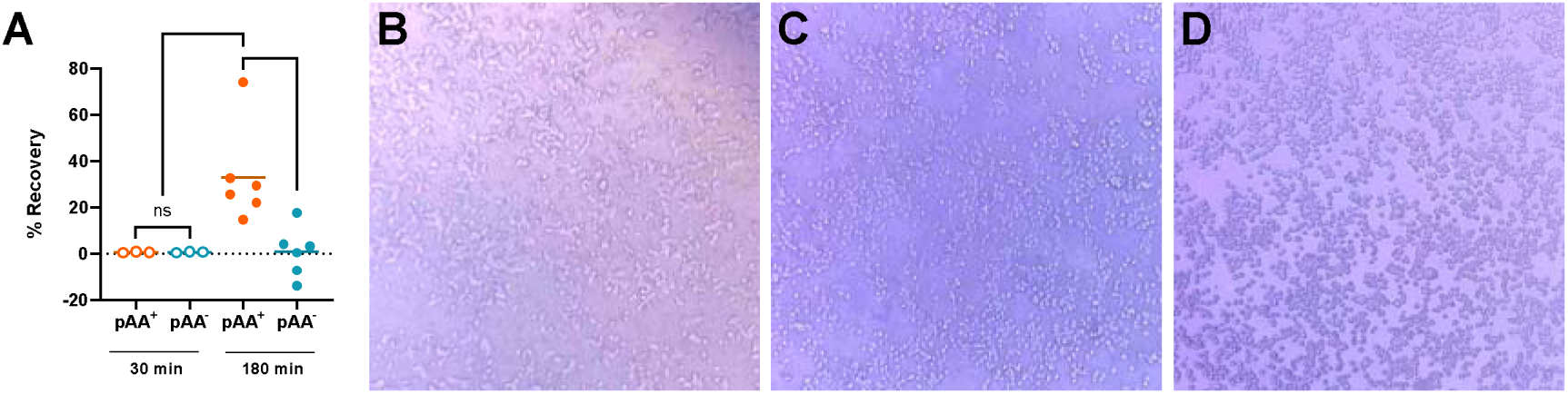
Effect of pAA on adherence of O104:H4 strain 1070/13 to HT-29 cells. **(A)** Per cent recovery of bacterial cells (normalised to inoculum) after incubation for either 30 min (empty circles) or 180 min (solid circles) with HT-29 cells. Each point is one biological replicate. Solid lines show the mean. Statistical analysis by one-way ANOVA with Tukey’s multiple comparison tests: ns, not significant (*P* ≥ 0.05); * *P* < 0.05; ** *P* < 0.01 (from *n* ≥ 3 biological replicates). **(B-D)** Light microscopy at 40x magnification of HT-29 cells at 180 min post-infection with either **(B)** 1070/13, **(C)** 1070/13 pAA^−^ or **(D)** no bacteria.

