## Supplementary Table for "Genetic and phenotypic analysis of the virulence plasmid of a non-Shigatoxigenic enteroaggregative *Escherichia coli* O104:H4 outbreak strain"

**Supplementary Table 1: Bacterial strains used in this study.**

| **Nomenclature** | **Species/serotype/genotype** | **Source** |
| --- | --- | --- |
| 1070/13 | Enteroaggregative *E. coli* O104:H4 strain 1070/13 | [28] |
| 1070/13 Δ*aggR* | EAEC 1070/13, Δ*aggR*::*cat* | This study |
| 1070/13 Δ*ccdB* | EAEC 1070/13, Δ*ccdB*::*cat* | This study |
| 1070/13 pAA^–^ | Plasmid-free derivative of EAEC 1070/13 | This study |
| *ccdB* Survival 2 T1^R^ | *E. coli* CcdB-resistant cloning strain (genotype unknown) | Invitrogen |
| DB3.1 | *E. coli* CcdB-resistant cloning strain (*gyrA462*) | Invitrogen |
| MFD*pir* Δ*hsdR* | *E. coli* MFD*pir* Δ*hsdR;* conjugation donor | C. M. Tang (Oxford) |
| MG1655 | *E. coli* K-12 MG1655 wild type | [39] |
| PAO1 | *P. aeruginosa* PAO1 wild type | [59] |

**Supplementary Table 2: Plasmids used in this study.**

| **Plasmid ID** | **Construction/purpose** | **Source** |
| --- | --- | --- |
| pAggR | pGM101_neo_ containing entire *aggR* locus | This study |
| pBAD33 | Arabinose-inducible vector for toxicity assays | [47] |
| pBAD33::*ccdB*_110_ | pBAD33 encoding CcdB_110_ | This study |
| pBAD33::*ccdB*_101_ | pBAD33 encoding CcdB_101_ | This study |
| pBAD33::*ccdB*_M90T_ | pBAD33 encoding CcdB_M90T_ | This study |
| pBAD33::*relE1* | pBAD33 encoding RelE/ParE 1 | This study |
| pBAD33::*relE2* | pBAD33 encoding RelE/ParE 2 | This study |
| pBAD33::*vapC*_142_ | pBAD33 encoding VapC_142_ | This study |
| pBAD33::*vapC*_133_ | pBAD33 encoding VapC_133_ | This study |
| pCcdB | pGM101_neo_ containing entire *ccdAB* locus | This study |
| pCONJ5K | Mobilisable vector for mutagenesis | [29] |
| pGM101 | Promoterless vector for antitoxicity assays | [26] |
| pGM101_neo_ | Neomycin-resistant variant of pGM101 | This study |
| pGM101_neo_::*ccdA* | pGM101_neo_ containing *ccdA* plus its promoter | This study |
| pIB279 | *sacB*-Neo^R^ cassette for pSTAB vectors | [52] |
| pMW_O104 | pSTAB with FII/FIB replicons from pAA_1070_ | This study |
| pMW_O104::*ccdAB* | pMW_O104 containing entire *ccdAB* locus | This study |
| pUC19 | High-copy cloning vector | New England Biolabs |

**Supplementary Table 3: Oligonucleotide primers used in this study.**

| **Primer ID** | **Sequence (5’-3’)** | **Target** |
| --- | --- | --- |
| GMntu106 | gcgcatcccagagggacatcttatattccccagaacatcaggttaatg | *ccdAB* for pSTAB |
| GMntu107 | gtgatgggttaaaaaggatccggatctgccggaaatgg |  |
| GMntu112 | gcgcatcccagagggacatcgatcctttttaacccatcacatatacctgccgttcac | *sacB-NeoR* |
| GMntu113 | gttaaaaaggatcgatgtccctctgggatgcgctccggatgaatatgatgatc | pAA_1070_ replicon |
| GMntu114 | caggacgacgaggcttgcaccttcataatcggtagtgtatgctgtttttctgg |  |
| GMntu115 | gcatacactaccgattatgaaggtgcaagcctcgtcgtcctggccggaccacgctatctg | *sacB-NeoR* |
| GMntu132 | ccatttccggcagatccggatcctttttaacccatcacatatacctgccgttcac | pMW_O104 for *ccdAB* insertion |
| GMntu133 | ctggggaatataagatgtccctctgggatgcgctccggatgaatatgatgatc |  |
| MS123 | cgaagcggcatgcatttacg | pGM101 for complementation |
| MS124 | ccttcgcgcgcgaattgatc |  |
| MS125 | gatcaattcgcgcgcgaagggactgggctgcaattaag | *aggR* for complementation |
| MS126 | cgtaaatgcatgccgcttcgtcattggcttttaaaataagtc |  |
| MS127 | gatcaattcgcgcgcgaaggtttcctcagtgtggtacac | *ccdAB* for complementation |
| MS128 | cgtaaatgcatgccgcttcgttatattccccagaacatcag |  |
| RWntu011 | ccgggtaccgagctcgaattc | pBAD33 |
| RWntu012 | ggatcctctagagtcgac |  |
| RWntu013 | aattcgagctcggtacccggcgacgaaggaagatttgac | *vapC*_142_ |
| RWntu014 | aggtcgactctagaggatccttacttcacccagtcttc |  |
| RWntu015 | aattcgagctcggtacccgggacccgaacgaagacctatatg | *vapC*_133_ |
| RWntu016 | aggtcgactctagaggatccttacttcacccagtcttcc |  |
| RWntu017 | aattcgagctcggtacccggttattgaaatgaacggctc | *ccdB*_110_ |
| RWntu018 | aggtcgactctagaggatccttatattccccagaacatcag |  |
| RWntu028 | agaaacgcaaaaaggccatc | pGM101 (to make pGM101_neo_) |
| RWntu029 | ctgtcagaccaagtttactc |  |
| RWntu030 | gatggcctttttgcgtttctcttcacgctgccgcaagc | *neoR* |
| RWntu031 | gagtaaacttggtctgacagaggcggcggtggaatcga |  |
| RWntu040 | tcgcgcgcgaaggcggatctgccggaaatgg | *ccdA* |
| RWntu041 | gcatgccgcttcgtcaccagtccctgttctc |  |
| RWntu074 | aggtcgactctagaggatccttatattccccagaacatcagg | *ccdB*_101_ |
| RWntu075 | aattcgagctcggtacccggttgccgatgagaacaggg |  |
| RWntu081 | aattcgagctcggtacccggaaatgaacggctcttttg | *ccdB*_M90T_ |
| RWntu082 | aggtcgactctagaggatccttatattccccagaacatcag |  |
| RWntu088 | aattcgagctcggtacccggactgacaatccgttaccg | *relE/parE 1* |
| RWntu089 | aggtcgactctagaggatccctacctgttttctcgtgtg |  |
| RWntu090 | aattcgagctcggtacccggaagctggcgcgagggctt | *relE/parE 2* |
| RWntu091 | aggtcgactctagaggatcctcagttgaaatgacggctggcatc |  |
| RWntu106 | tcgtttgaaatcccgcttcg | *dnaA* (ddPCR) |
| RWntu107 | cgttttcgtcggcctttttc |  |
| RWntu108 | agccgatccgttaaaagcac | *fliC* (ddPCR) |
| RWntu109 | tggtggtgttgttcaggttg |  |
| RWntu110 | tgttaacgaacgtcgcgaag | *hns* (ddPCR) |
| RWntu111 | attgctgcagcttacgagtg |  |
| RWntu112 | gctgtaggacccacttattagc | *aggA* (ddPCR) |
| RWntu113 | tcctccacaaactgttggtg |  |
| RWntu114 | atatcggtggtcatcatgcg | *ccdB* (ddPCR) |
| RWntu115 | aagtctcccgtgaactttaccc |  |
| RWntu116 | gcaatagccaatttgcacat | *wzy* (ddPCR) |
| RWntu117 | cccggggcaattatcattaa |  |
| RWntu140 | ggtacccggggatcctctag | pUC19 |
| RWntu141 | gagctcgaattcactggcc |  |
| RWntu142 | cggccagtgaattcgagctccagtgttgcaaaaatggtg | *aggR* downstream |
| RWntu143 | cagcctacacttgacttattttaaaagccaatg |  |

**Supplementary Table 3 cont.: Oligonucleotide primers used in this study.**

| **Primer ID** | **Sequence (5’-3’)** | **Target** |
| --- | --- | --- |
| RWntu144 | aataagtcaagtgtaggctggagctgcttc | *cat* (Δ*aggR*) |
| RWntu145 | tgataaagacatgggaattagccatggtcc |  |
| RWntu146 | taattcccatgtctttatcaggcaactctgc | *aggR* upstream |
| RWntu147 | ctagaggatccccgggtaccggcgcgcatcctgtatattattg |  |
| RWntu148 | cagcctacacctgatgttctggggaatataaatgtc | *ccdB* downstream |
| RWntu149 | cggccagtgaattcgagctcgcgctttctctgtcctgc |  |
| RWntu150 | tggtgaaatgatgggaattagccatggtcc | *cat* (Δ*ccdB*) |
| RWntu151 | agaacatcaggtgtaggctggagctgcttc |  |
| RWntu152 | ctagaggatccccgggtaccctccctgacctgtgatgc | *ccdB* upstream |
| RWntu153 | taattcccatcatttcaccagtccctgttc |  |
| RWntu154 | aacgacggccagtgtttaaaggcgcgcatcctgtatattattg | *aggR* (deletion) |
| RWntu155 | agcaggaaacagctatgacgcagtgttgcaaaaatggtgg |  |
| RWntu156 | cgtcatagctgtttcctg | pCONJ5K |
| RWntu157 | tttaaacactggccgtcg |  |
| RWntu158 | agcaggaaacagctatgacggcgctttctctgtcctgc | *ccdB* (deletion) |
| RWntu159 | aacgacggccagtgtttaaactccctgacctgtgatgc |  |

**Supplementary Table 4: ddPCR master mix.**

| **Component** | **Concentration or quantity per 20 μl reaction** |
| --- | --- |
| 2x ddPCR Supermix for Probes (No dUTP) | 10 µl |
| Target primer/probe mix (FAM) | 900 nM primers/250 nM probes |
| Reference primer/probe mix (HEX) | 900 nM primers/250 nM probes |
| DNA | 50 ng |
