## Supplementary figures and images for "Genetic and phenotypic analysis of the virulence plasmid of a non-Shigatoxigenic enteroaggregative *Escherichia coli* O104:H4 outbreak strain"

### Supplementary Figure 1

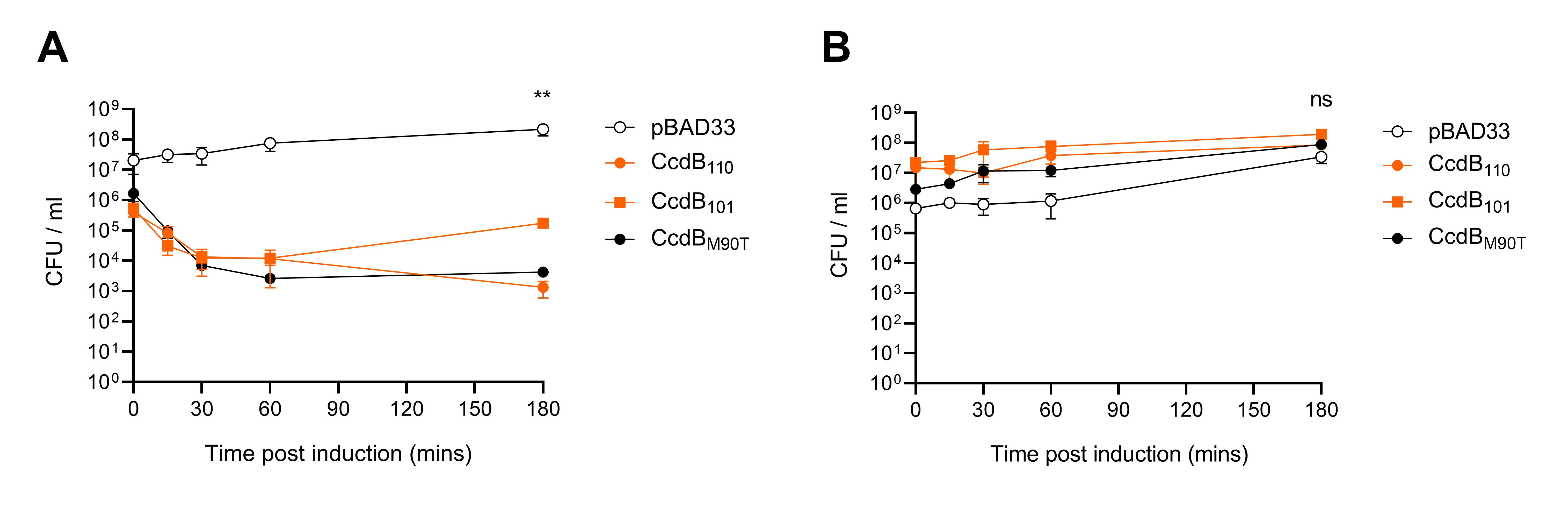

### Supplementary Figure 2

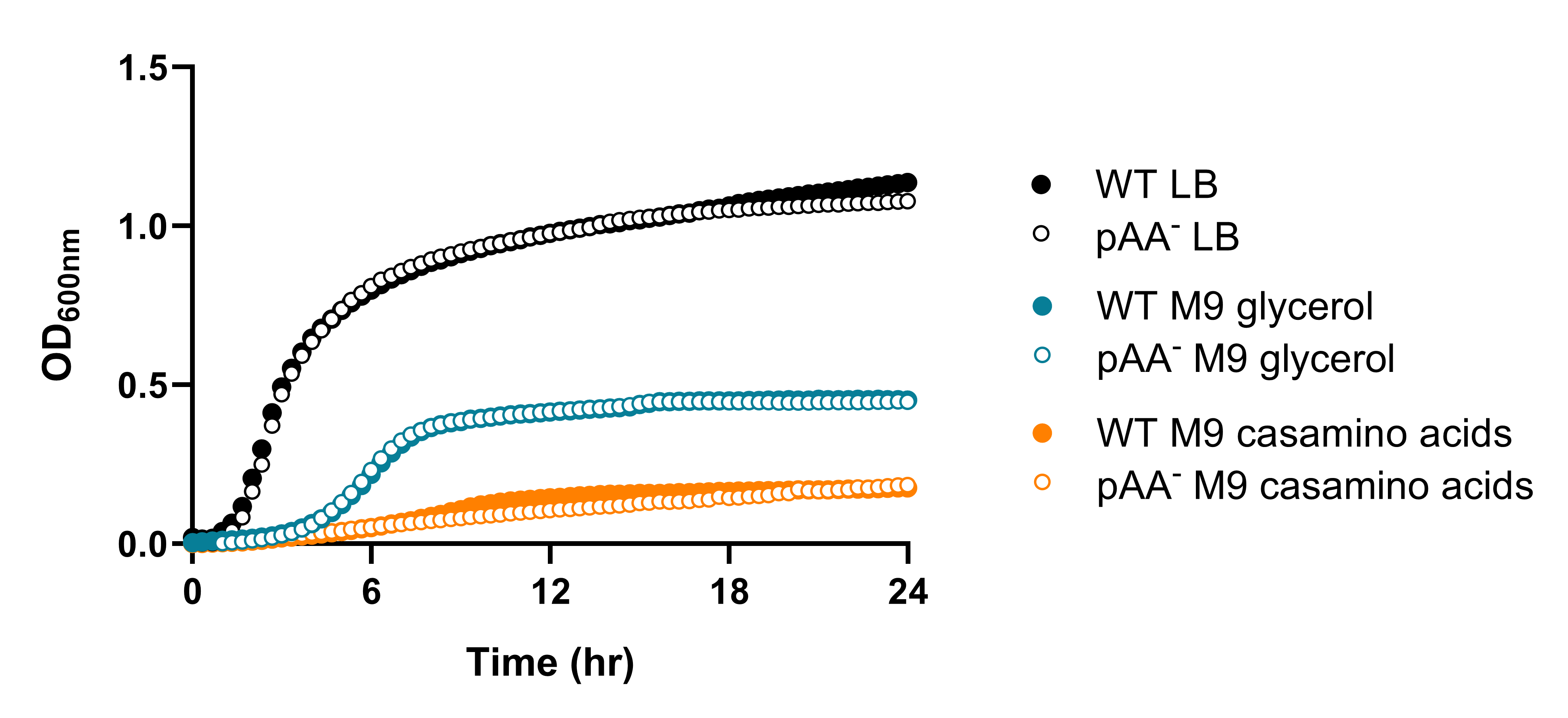

### Supplementary Figure 3

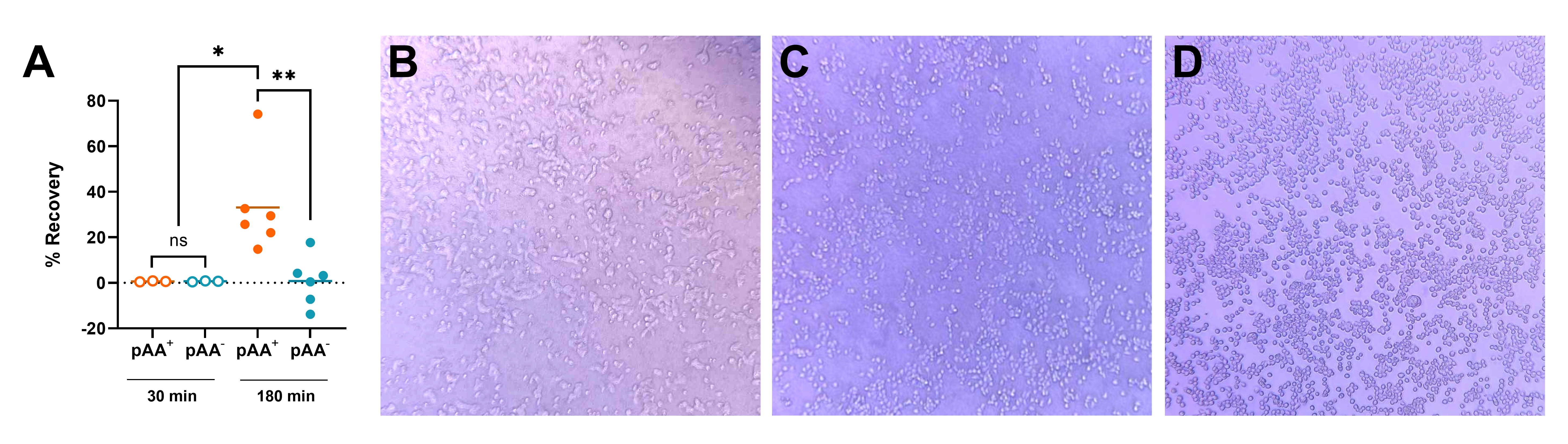
